# Single-molecule Mapping of Amyloid-β Oligomer Insertion into Lipid Bilayers

**DOI:** 10.1101/2024.02.21.581487

**Authors:** Arpan Dey, Abhsihek Patil, Senthil Arumugam, Sudipta Maiti

## Abstract

The interaction of disease-causing amyloid oligomers with lipid membranes is implicated in their toxicity. However, understanding the membrane interaction of different oligomers, and each constituent monomer in a given oligomer, has remained a challenge. Here we employed a recently developed single-molecule technique, called QSLIP, which can simultaneously determine the stoichiometry and membrane location of individual fluorescent labels on oligomeric membrane proteins. Using QSLIP, we measured the membrane insertion of small amyloid-beta (Aβ) oligomers of three different isoforms at the single-molecule level, and found that their toxicity is correlated with the depth of penetration of their amino-terminal into the bilayer. Such single-molecule maps provide a detailed assay for measuring the effect of any drug candidate on oligomer-membrane interactions.

## Main Text

In Alzheimer’s disease (AD), oligomers of Aβ peptides are the primary molecular agents responsible for neuronal damage^1–3^. Although a mechanistic picture of toxicity is yet to fully emerge, one of the most likely pathways involves their interaction with lipid bilayer membranes^4–6^. Such interactions may lead to membrane disruption^7^ or pore formation^8,9^, potentially resulting in cell death. The membrane interaction of amyloid oligomers has been studied for decades. However, the major question remains: Which specific oligomer is the most toxic, and why?

We have previously shown important structural features of Aβ oligomers by trapping oligomers during aggregation (flash-freezing) and subsequent solid state NMR^10,11^. Similarly, fluorescence and Raman based experiments of Aβ40 on lipid-bilayer coated nanoparticles established the antiparallel β sheet conformation of Aβ40 oligomers in membranes, with a C-terminus more buried than the N^12,13^. Structure of Aβ oligomers have also been studied using EPR (both radical labelled Aβ^14,15^). Another possibility is the usage of radical labelled lipids in membrane^16,17^. However, these studies did not have oligomer-wise resolution and therefore did not provide any correlation between stoichiometry, structure and toxicity.

Electrical recordings both *in vitro* and *in vivo* have shown that Aβ (both the 40 and 42 variants) can form pore-like structures in membranes^8,18–20^. Subsequent molecular modelling of these β-sheet rich pore-forming oligomers (PFOs) has also been performed^21,22^. Some structural information on β-PFOs by Aβ42 in micelles have been reported using NMR studies^21^. Subsequently, by using chemical cross-linking, SDS-PAGE, and MALDI-TOF, the same group also succeeded in identifying the stoichiometry (Aβ42 tetramers) of these β-PFOs^23^. This study reported the structure of a β-PFO, though using a non-physiological pH, and with micelle stabilised oligomers. In addition, a majority of these experimental observations are carried out under non-physiological conditions, e.g. at high concentrations (∼mM)^17,24^, which contrasts with the physiological plasma concentrations of Aβ (∼nM)^25,26^. At such concentrations, Aβ solution contains not only monomers and small oligomers, but also large oligomers, protofibrils, and fibrils^27^. Although isolation of intermediates (oligomers, protofibrils, fibrils) using techniques such as size exclusion chromatography (SEC) has been attempted^28^, identifying and differentiating individual small soluble oligomers has remained a challenge. The dynamic equilibrium between the small oligomers is a major complicating factor in characterizing individual oligomeric species. Most studies dealing with oligomers cannot differentiate between oligomers of different sizes, hence they average over the properties of individual species. We note that Aβ monomers can be separated from the oligomers, and have been shown to have a much lower affinity to lipid membranes and lower toxicity than the oligomers (dimers and larger) ^29,30^. However, stably separating each oligomer in a physiological buffer has not been possible. Single molecule techniques such as interferometric scattering (iSCAT) provides label-free approach in determining oligomerization, dynamics, and even structure of proteins^31,32^. Recently, a native mass spectrometric (NMS) study has reported the determination of conformational arrangement of detergent-solubilized membrane proteins reconstituted into membrane-mimetic liposomes^33^. However, the structure of individual Aβ oligomers in membrane under physiological conditions is still beyond reach. Therefore, determining molecular details of how individual types of Aβ oligomers interact with the membrane, and whether that correlates with their toxicity, remains a major unsolved problem in deciphering Alzheimer’s disease mechanism.

The problem can be separated into two independent parts. First, a lack of techniques to identify and study the membrane interaction of each type (i.e. stoichiometry) of Aβ oligomers among a pool of many. Second, a lack of proper experimental strategy to correlate the membrane interaction with the toxicity of each type of oligomer. Here we addressed the first problem with a newly devised single-molecule technique called Q-SLIP, which allows the simultaneous determination of the stoichiometry of each individual oligomer in a mixture, and the surface exposure of each individual monomer in each type of oligomer^34^ (*vide infra*). Next to address the second problem, we adopted an indirect approach. We measured differences in the membrane interaction of three different Aβ isoforms with known difference in toxicity. We probed small oligomers of three Aβ isoforms, namely Aβ42, Aβ40, and AβCha (a synthetic isoform with F19 replaced by cyclohexylalanine^35^, complete sequences of the peptides are given in SI 1). It is known that Aβ42 oligomers are more toxic than Aβ40^36^, and AβCha (with the crucial F19-L34 contact disturbed^35^) is the least toxic. If toxicity is dependent on interaction with membrane interaction, then different oligomers of these different Aβ isoforms should interact differently with the membrane.

Isoform-specific differences in putative membrane-assisted toxicity have one of three likely origins. There can be differences in 1) the nature of oligomers formed by each isoform in the solution state, prior to membrane attachment, 2) the relative membrane affinity of these oligomers, or 3) the nature of insertion and the conformation adopted by each type of oligomer inside the lipid bilayer. These three can also operate in any combination. Here we test all three possibilities, and then ask if any observed difference correlates with the relative toxicity of the species.

Each oligomeric species formed in the solution state can be probed by single molecule photobleaching^37–39^ (smPB). Stoichiometry of fluorescently labelled Aβ aggregates can be identified by counting the steps in their corresponding bleaching trajectories^37,40^. The relative membrane affinity of individual oligomers (termed as relative affinity) can also be measured using smPB^37^. However, smPB alone cannot provide any information about conformation or membrane penetration of the oligomers.

This problem can be tackled by the recently developed method of Quencher Induced Step Length Increase in Photobleaching (QSLIP)^34^. Processes such as dynamic quenching and labile complexation with a quencher can impart photostability to a fluorophore, since quenching reduces its probability to be in the singlet or triplet excited state from where photobleaching typically occurs^41^. QSLIP measures this photostabilization (i.e. an increase in time that a fluorophore takes to bleach), and therefore it reports the accessibility of a fluorophore to an external quencher. If a quencher (in our case, a doxyl radical labeling a lipid chain) is placed at a specific location in a membrane, e.g. at the 5’ or 16’ positions, and can quench a fluorophore on contact (in our case, a rhodamine-B labeling the amino terminal of each Aβ monomer), then QSLIP would yield the relative proximity of the amino terminal to these two locations in the membrane. Also, since this method is based on the photobleaching probability of individual monomers, it not only automatically provides the stoichiometry of each oligomer, but also reports the accessibility of each individual monomer in the oligomer. Thus, QSLIP provides a detailed measure of the membrane penetration of each monomer in each type of Aβ oligomer.

To verify that the differences of toxicity between the three species, we measured the reactive oxygen species (ROS) induced by their oligomers in a rat neuronal cell line RN46A. ROS reports cellular stress, and is therefore linked to toxicity^42,43^. We incubated RN46A in 3 μM Aβ for 30 minutes (experimental detail in SI 2). We note that the higher concentration (compared to other experiments reported here) was used to unequivocally determine the relative levels of Aβ toxicity within a reasonable experimental time. The ROS levels were reported by the fluorescent reporter dye CellROX Green. As shown in **Figures 1A, 1B**, and **1C**, maximum ROS production was observed for Aβ42, followed by Aβ40, and then AβCha. The results indicate that cell toxicity (in terms of ROS generation) agrees with previously established results, and follows the order Aβ42 > Aβ40 > AβCha.

**Figure 1:**
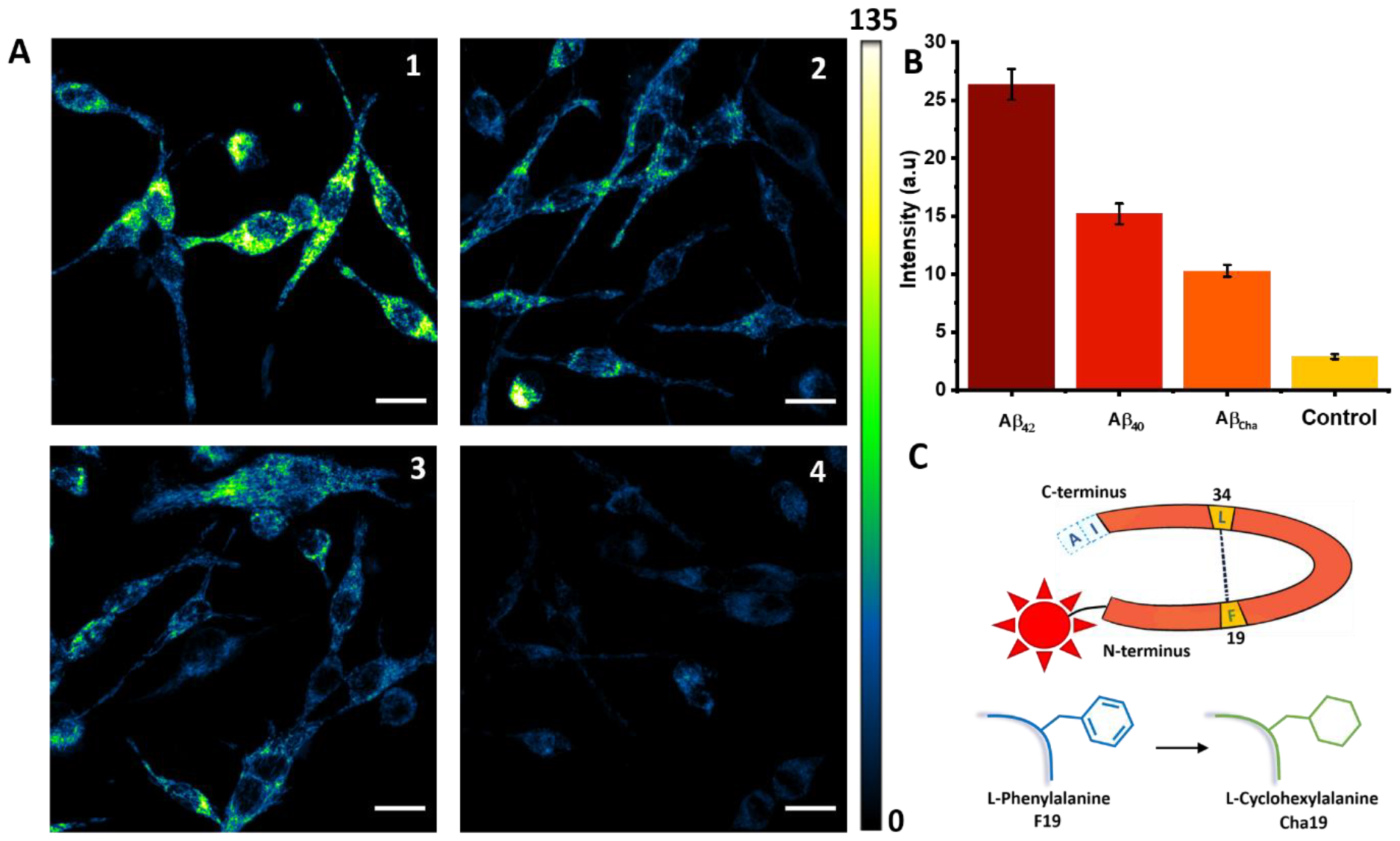
ROS measurements in RN46A cells. (A) Confocal image of cellROX™ green intensity from RN46A cells upon incubation with 3μM of Aβ oligomers. A1, Aβ42; A2, Aβ40; A3, AβCha, and A4, control (i.e. no added Aβ). Scale bar = 20 μm. (B) Average ROS intensity obtained from the cells (mean ± SE, n=3 independent measurements). (C) A cartoon representation of the three different Aβ variants. N-terminus of the peptides were labelled with rhodamine B. Orange part depicts Aβ40, the contact between F19 and L34 (both yellow) is marked by a dashed line. Aβ42 has two additional amino acids I and A, at the C-terminus. AβCha has F19 mutated to cyclohexylalanine which abolishes the contact (Cha, lower part).

We then tested the first of the likely origins of the differences in toxicity between the three isoforms of Aβ. We measured whether the size distribution of the oligomers in solution is different between the three species using single molecule photobleaching (smPB). The details of the experiment are given in SI 3. Briefly, N-terminus rhodamine B-labelled Aβ (RAβ) oligomers were diluted in Thomson’s buffer (TB) from high pH stocks (pH 11). The resulting solution was diluted with a 0.25% polyvinyl alcohol (PVA) solution (W/V) to a sub-nanomolar concentration, and this was spin-coated onto pre-cleaned coverslips. Many regions of each coverslip were imaged using a home-built Total Internal Reflection Fluorescence (TIRF) microscope (fig S1). A representative Region of Interest [ROI] from one of the images is shown in fig S2A. The photobleaching steps were identified by a Napari-based step detection algorithm developed by us (typical representative traces and fits are given in S2 B, S3), and many of them were also checked manually. The number of bleaching steps obtained from a given fluorescent spot revealed the stoichiometry of the oligomer present at that spot. The concentration was low enough so that the probability of having two oligomers occurring within the same spot (whose size is determined by the optical resolution limit) was negligible. The solution state oligomer distribution was monomer-heavy, and was similar among all three Aβ isoforms (Aβ 42, 40, and Cha, as shown in **Figure 2A**). However, the relative amount of dimers compared to the monomers was somewhat higher for RAβ42 (38.8% monomer and 34.4% dimer) than the others, but the difference was not statistically significant. It is important to note that these distributions were not pre-bleaching corrected (pre-bleaching is an experimental artifact^44^). So the absolute distributions may be somewhat skewed towards smaller oligomers, but relative differences in population across isoforms should be robust even without this correction. So we infer that the difference between the oligomer size distribution between the three isoforms in the solution state does not explain the observed differences in their toxicities.

**Figure 2:**
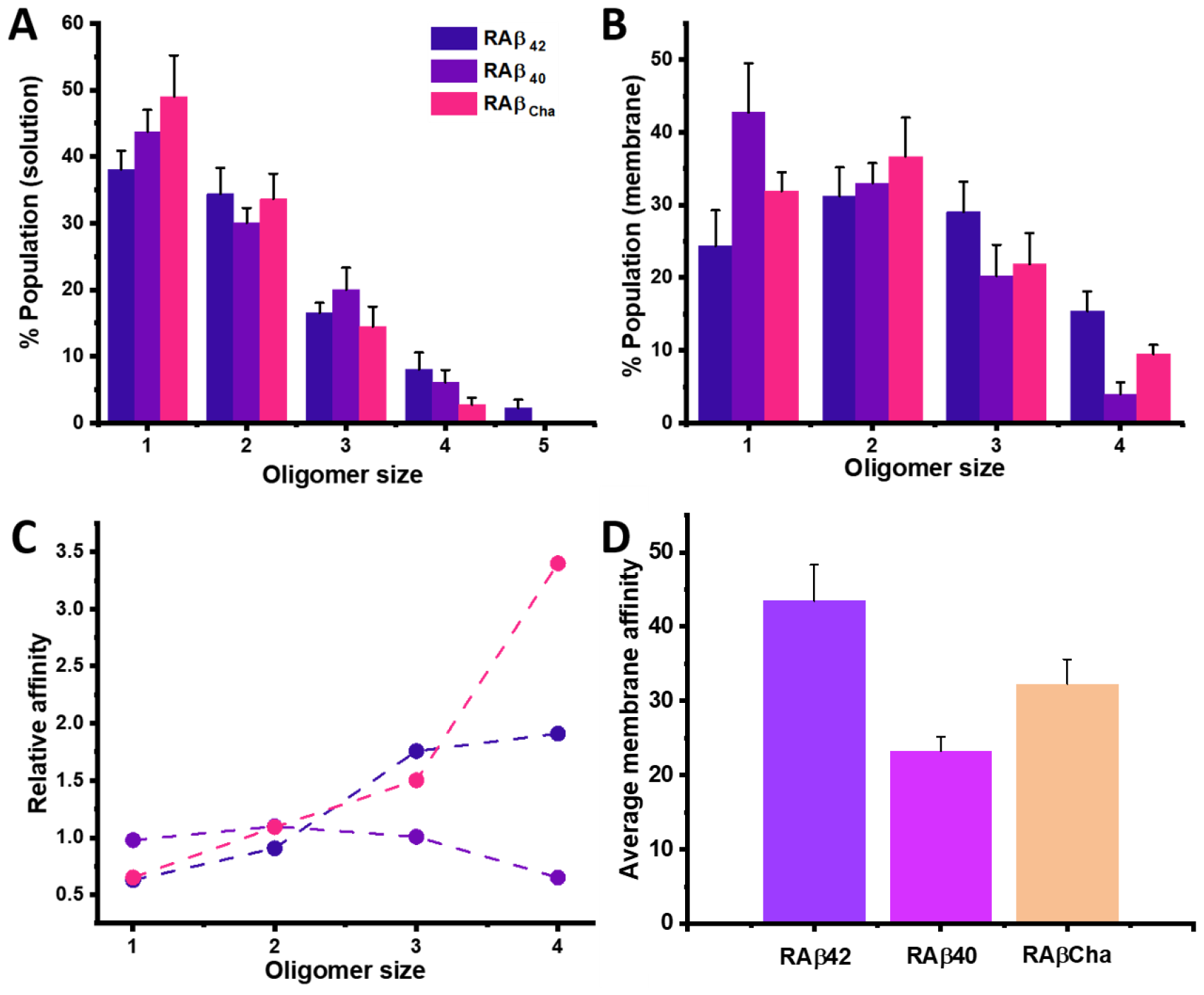
Relative membrane affinity of individual oligomeric species of different Aβ isoforms. (A) Size distribution in solution (Aβ42, blue, n=5; Aβ40, purple, n=9, and AβCha, magenta, n=5). (B) Size distribution in the membrane (n>=5). (C) Relative membrane affinity of individual oligomers of different Aβ variants. Values in A and B are mean ± SEM. Colour scheme is the same throughout. (D) The overall membrane affinity averaged over oligomers. Values represent mean ± SEM, n>=3.

The second possibility was that the membrane affinity of individual oligomers of the three Aβ isoforms are different. We measured the stoichiometry of RAβ oligomers attached to a supported lipid bilayer (POPC:POPG:Cholesterol in a molar ratio of 1:1:1; details of bilayer preparation are given in SI 4) using smPB. Oligomers were incubated on the bilayer for 30 minutes, then washed to remove excess unattached oligomers, and finally imaged under TIRF illumination until the oligomers bleached completely. The membrane distribution of RAβ oligomers is shown in **Figure 2B**. We calculated the relative affinity of individual oligomers for PPC 111 membrane by comparing the stoichiometric distribution in the membrane and in the solution. For a specific size oligomer (say an n-mer), it can be calculated as

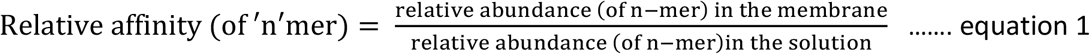

As shown in **Figure 2C**, the trimer and tetramer of RAβ42 have higher membrane affinity (1.76 and 1.91, respectively) compared to the monomer or the dimer (0.63 and 0.91, respectively).

However, for RAβ40, the relative affinities of the monomer, dimer, and trimer were comparable (0.97, 1.09, and 1.0, respectively), and these were higher than that of the tetramer (0.65). For the less toxic RAβCha, the relative affinity showed a gradual increase for the larger oligomers (0.65, 1.09, 1.50, and 3.39 for the monomer, dimer, trimer, and tetramer, respectively). The average membrane-affinity of each species (that is, the total number of oligomers attached to an area of 31.2×31.2 μm^2^ in a bilayer for the same concentration of Aβ incubation) yielded the order RAβ42 > RAβCha > RAβ40 (as shown in **Figure 2D**). This order does not correlate with their observed toxicity too. Thus, collectively, neither oligomer stoichiometry in the solution state, nor their membrane affinity could explain the differences in toxicity between the different Aβ isoforms.

Therefore, if the membrane interaction deteremines toxicity, the only remaining possibility is that the difference lies in how each type of oligomer inserts into the lipid bilayer membrane. We performed QSLIP measurements to determine the membrane location of each monomeric constituent of each oligomeric species. In the photobleaching trajectories (fig S2B) the time taken for an individual fluorophore to bleach (t = 0 is when the excitation is turned on) is referred to as the ‘step length’. A ratio of such step lengths in presence of a quencher to that in its absence provides information on the extent of quenching-induced stabilization. This is a measure of the accessibility of the fluorophore to a quencher. Thus the QSLIP value is defined as

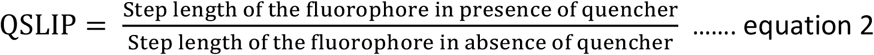

Since quenching is expected to increase the longevity of the fluorophore by reducing the time spent in the triplet state, a value much greater than 1 indicates close proximity of the quencher to the fluorophore. Values near unity indicate a fluorophore shielded from the quencher.

We used quenchers that can interact with the fluorophore (the N-terminus rhodamine label of each peptide) at three different depths of the bilayer. Tryptophan is a water-soluble quencher, and when added to the solution, can measure the solvent exposure of the membrane-attached oligomers. However, if the N-terminus is substantially buried into the bilayer, then tryptophan quenching cannot determine the extent of bilayer penetration. On the other hand, doxyl radicals anchored at different positions of the lipid fatty acyl tail can probe the N-terminus location of the oligomers inside the bilayer, as they are effective contact-based quenchers of the triplet state of rhodamine^45^. Here we used 3 mol % doxyl anchored lipids 16:0-16 doxyl PC (doxyl labelled at the 16^th^ carbon of the stearyl tail, abbreviated as C-16 doxyl henceforth) and 16:0-5 doxyl PC (abbreviated as C-5 doxyl). Since the radicals were positioned at different depths of the bilayer, QSLIP values influenced by these radicals provided a good measure of the location of the N-terminus inside the bilayer.

With our experimental protocol, we could obtain statistically reliable results only till the tetramer, as there were very few larger oligomers. The bleaching events in the bleaching trajectories were sequentially labelled as step 1, step 2, step 3, and step 4 respectively (as shown in fig S2B). Monomers only had step 1, while a tetramer had all the 4 steps. The monomer step lengths of RAβ42, RAβ40, and RAβCha are provided in **figure 3A, B**, and **C** respectively (all the oligomer step lengths are provided in table 1 in the SI). It can be clearly seen that under the different quencher conditions, the RAβ42 monomer shows the highest increase in the step length (Q-SLIP value of 3.21 ± 0.62) for the C-16 doxyl labelled bilayer. This suggested that the N-terminus of the Aβ42 monomers penetrated deep into the bilayer, almost to the mid-plane. However, none of the other two monomers showed such high QSLIP values, indicating a lack of preference for any specific location in the bilayer for them. This suggested a difference among the relative positioning of the N-terminus of different Aβ monomers in a bilayer (other oligomers are mentioned subsequently, data in table 1, SI).

**Figure 3:**
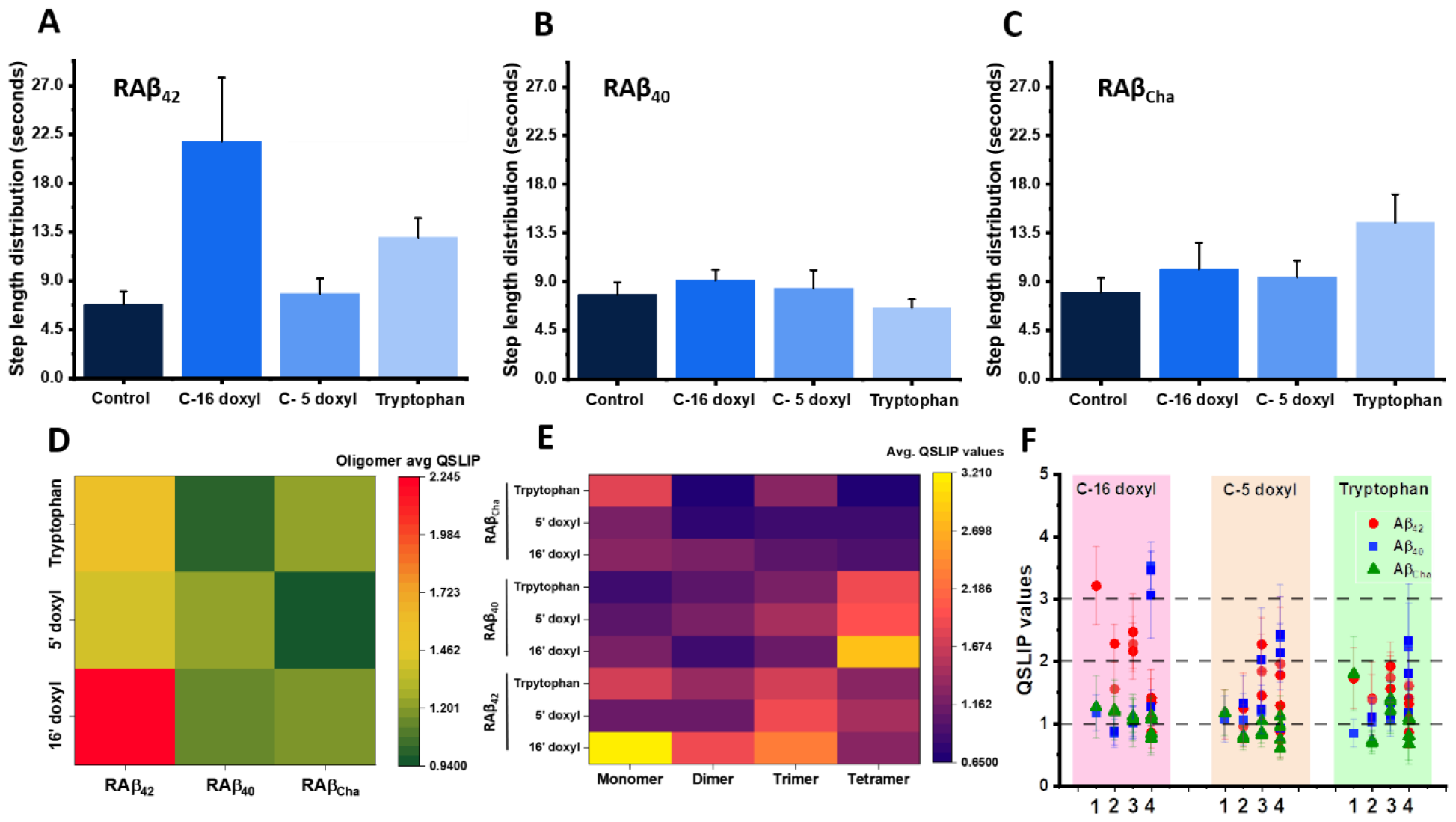
Step length distributions of the monomer of different Aβ isoforms in PPC111 bilayer with different quenchers (A) Step length of Aβ42 monomer, (B) Aβ40 monomer, and (C) AβCha monomer. Values are mean ± SEM. Colour code is the same for all the plots. (D) Amplitude weighted QSLIP values of all the oligomers with C-16 doxyl, C-5 doxyl, and tryptophan. (E) QSLIP values of oligomers with the three different quenchers. Here, the QSLIP values of all the monomers in an oligomer were averaged, and (F) the QSLIP values of each of the individual monomers in an oligomer. Data is mean ± SEM. The x-axis represents the stoichiometry of the corresponding oligomers.

As seen in equation 2, QSLIP normalizes the step lengths of individual RAβ oligomers by the step length in absence of any quencher. It therefore takes into account any isoform-specific variation in the bleaching probability. Thus one can compare the QSLIP values across different Aβ isoforms. The membrane penetration of each Aβ isoform is represented by the amplitude weighted average QSLIP values over all the oligomers. This is a product of the oligomer distribution of an Aβ isoform and the QSLIP value of individual Aβ oligomers. It is defined as

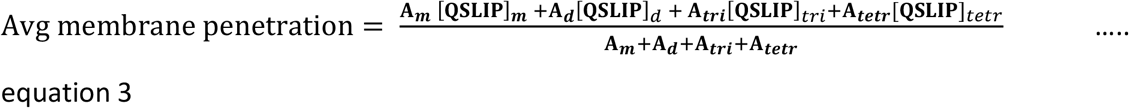

Where **A**_**n**_’s are the average population of an oligomer in the membrane, and [QSLIP]_n_ is the corresponding QSLIP value (*m* = monomer, *d* = dimer, *tri* = trimer, and *tetr* = tetramer). Amplitude averaged QSLIP values of the three Aβ isoforms showed different penetration abilities. The N-termini of Aβ42 oligomers preferentially gets buried deeper in the bilayer and reach near the C-16 position of the lipid tail (with values of 2.24 for C-16 position, 1.39 for C-5 position, and 1.56 for solvent exposed). Aβ40 on the other hand show a mild preference for the C-5 position (values of 1.11 for C-16 position, 1.23 for C-5 position, and 1.02 for solvent exposed). Compared to the other two isoforms, the N-terminus of AβCha remained solvent exposed without much penetration (values of 1.12 for C-16 position, 0.95 for C-5 position, and 1.22 for solvent exposure). A heat map of these average QSLIP values across all oligomers is provided in **figure 3D**. The results also show that membrane penetration by Aβ42 is unmatched by all the other isoforms (marked by the high QSLIP value with C-16 position). Although amplitude averaged QSLIP of Aβ can also be obtained from bulk measurements^21^, additional information can also be obtained from it. Since it is based on a single-molecule photobleaching technique, it can resolve individual bleaching events, and leads to the identification of oligomers. **Figure 3E** shows a heat map of average QSLIP values from individual oligomers of the three Aβ isoforms under three different quencher concentration. This is obtained by averaging the QSLIP of individual subunits in an oligomer. A gradual decrease in the average QSLIP values can be seen moving from RAβ42 to RAβCha. This suggested a membrane-penetration ability in the order Aβ42 > Aβ40 > AβCha.

These results show that the membrane location of different isoforms of Aβ correlates with their toxicity. We do not know whether this was a direct consequence of difference in the structure adopted by these different oligomers, but we note that the N-termini of all three oligomers are identical upto residue number 18. As already mentioned, QSLIP provides information on the individual bleaching steps within an oligomer. These contains membrane location information of the individual monomers which constitute an oligomer. The QSLIP values of individual subunits of the three Aβ isoforms are provided in **figure 3F** (values in table 2, SI). In Aβ42, the general trend is for the N-terminus to insert deep till the C-16 position (with a high QSLIP value especially for the monomers). However, the two steps indicate different QSLIP values for the dimer near C-16 (2.28 ± 0.31, and 1.55 ± 0.39). This indicated that the N-termini of the two monomers in an Aβ42 dimer are not located in a similar place. The absence of a prominent difference in QSLIP values along with low absolute values indicated that the N-termini of the dimer remains far away from them. On the contrary, the trimer adopts an in-register conformation with similar QSLIP values for all the three subunits. A higher absolute value for the C-16 position over other positions indicates a preferred localization near the membrane center. The tetramer however behaves completely differently with QSLIP value nearly unity for all the quenchers. This indicates that the hidden N-termini of the tetramer are relatively unexposed to the membrane. In case of Aβ40, most of the oligomers and subunits attain a geometry similar to the tetramer of Aβ42, that is, compact and unaffected by any of the quenchers (blue squares in **figure 3F** including the standard error). However, the tetramer stands out in terms of its conformation with three of its subunit s comparatively exposed and one compact. The absolute values are higher for the C-16 position.

Contrasted against specific oligomers of different conformations (trimer and tetramer) of Aβ42 and 40, all the oligomers of AβCha are highly inaccessible to all the quenchers. This indicated a compact conformation/position attained by these oligomers that also likely prevented membrane penetration.

From our QSLIP measurements, the highest QSLIP values were obtained from the monomer of RAβ42 and the tetramer of RAβ40. We further corroborated these high QSLIP values by varying the percentages of radical-labelled lipids in a bilayer and probed the ‘dose dependence’ of the quencher radical (SI6, fig S4). The results are consistent with previous observations of deeper membrane penetration by RAβ42 compared to RAβ40.

Understanding the structure of small Aβ oligomers experimentally has been a major challenge. A structured membrane pore, to be toxic, likely needs to have specific locations for its constituent parts and span the entire width of a bilayer. This is what we aimed to measure here. Our results demonstrate that the oligomers of Aβ42 and Aβ40 indeed have reasonably well-defined strutures with specific locations inside the membrane. Moreover, the pattern of insertion of each of the monomers of an oligomer correlates with the difference in toxicity among the small oligomers of Aβ42 and Aβ40. Under similar incubation times, the N-terminus of Aβ42 reaches closer to the centre of the bilayer (near C-16), whereas Aβ40 extends mostly to the C-5 position. AβCha, the least toxic species, has a more folded conformation than either. Thus, the greater penetration of the amino-terminal of Aβ oligomers correlates with higher toxicity. We note that there is more detailed information provided by QSLIP than this simplifying description of their differences. This information can in principle be utilized to examine the effect of candidate drug molecules on the oligomers. It can also provide valuable constraints for constructing a model of Aβ oligomer structure in the membrane. Our result demonstrates the general applicability of QSLIP in mapping the intricate structural organization of membrane-incorportaed protein oligomers at the single-molecule level.

## Author Contributions

AD performed all the experiments, AP and SA developed the analysis routine for automated step detection and helped in analysing the data, AD and SM conceptualized the project and co-wrote the paper.

## Acknowledgment

This research has been supported by a Dept. of Atomic Energy, Govt. of India, intramural grant to SM (no. RTI 4003).

