## Supporting information for "Single-molecule Mapping of Amyloid-β Oligomer Insertion into Lipid Bilayers"

### Table of contents

**SI 1.** **Sequence of Aβ (42, 40, and F19Cha)**

**SI 2.** **Standardization of Aβ induced ROS measurements in RN46A cells**

**SI 3.** **Single molecule photobleaching (smPB) – experimental details, and step detection**

**SI 4.** **Preparation of supported lipid bilayer**

**SI 5.** **QSLIP values for all RAβ variants**

**SI 6.** **Doxyl dose-dependent change in step length of RAβ42 vs RAβ40 oligomers**

**SI 1: Sequence of Aβ (42, 40, and F19Cha)**

The sequence of Aβ peptides used for our experiments are as follows:

Aβ42 – DAEFRHDSGYEVHHQKLVFFAEDVGSNKGAIIGLMVGGVVIA

Aβ40 – DAEFRHDSGYEVHHQKLVFFAEDVGSNKGAIIGLMVGGVV

Aβ(F19Cha) – DAEFRHDSGYEVHHQKLV*(Cha)*FAEDVGSNKGAIIGLMVGGVV

**SI 2: Standardization of Aβ induced ROS measurements in RN46A cells**

We measured the reactive oxygen species (ROS) in RN46A cells to estimate the intracellular toxicity of the Aβ oligomers. 3 µM unlabelled Aβ peptide (stock concentration 1 mM, pH 11) was used to estimate the cell ROS. Aβ was incubated to the cells in media for 30 mins. Post incubation, the excess Aβ was washed off thrice using TB. 5 µM cellROX$^{TM}$ Orange Reagent (Excitation: 633 nm, Emission: 641 nm – 700 nm) was incubated for another 30 mins. As a sham control, Aβ was replaced with buffer and incubated for the same duration. The cell ROS was quantified by monitoring the average fluorescence intensity from z-projections of the cells.

**SI 3: Single molecule photobleaching (smPB) – experimental details**.

To perform smPB of RAβ oligomers of dthe three Aβ isoforms, we used a home-built objective based total internal reflection fluorescence (TIRF) microscope. A 543-nm He-Ne (25-LGR-393-230; Melles Griot, Rochester, NY) laser was used for focused excitation (back aperture power was maintained at 1.3 mW) of a Nikon APO TIRF 100X/1.49 objective. A dichroic (565 nm) was used to separate the fluorescence from the excitation. The fluorescence was collected using a band-pass filter (605/55 nm, BA577-633, Nikon) and focused using a 50 cm biconvex achromatic doublet onto an electron multiplying CCD camera (ANDOR iXON, DV887ECS-UVB). A diagrammatic representation of the TIRF instrument is provided in **figure S1**.

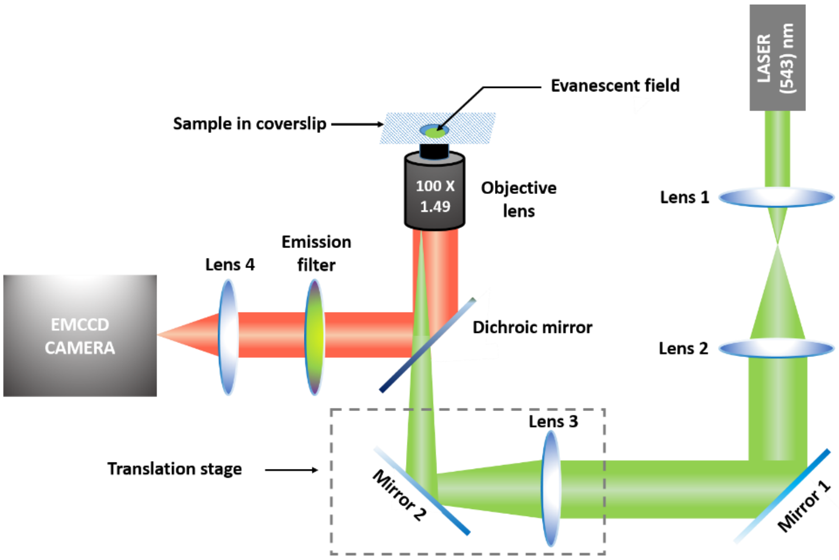

**Figure S1: Schematic representation of the home-built objective based TIRF setup. A 543 nm HeNe Laser is used as a source of excitation. The translation stage coupled with a lens and mirror is used to switch between TIR excitation and wide field excitation. The laser is focused at the back-focal plane of the objective to attain a geometry for TIR excitation. The emission is separated using a dichroic filter and collected and focused to an EMCCD camera using lens 4.**

In smPB, to detect the stoichiometry of oligomers, the fluorescent markers should be stationary. This allows artifact free detection of the bleaching steps. To detect the solution state oligomerization of the different RAβ isoforms, they were freshly prepared in pH 7.4 (from a stock solution of 30 µM, pH 11) in Thomson's buffer (NaCl - 146 mM, KCl - 5.4 mM, $CaCl_2.2H_20$ - 1.8 mM, $MgSO_4$ - 0.8 mM, $KH_2PO_4$ - 0.4 mM, $Na_2HPO_4$ – 0.3 mM, d-Glucose – 5 mM, Na Hepes – 20 Mm) and diluted in 0.25% poly vinyl alcohol (PVA) solution to prepare a final concentration of 0.5-1 nM. The resulting PVA solution with the oligomers were spin-coated on coverslips (precleaned, treated with piranha solution and cleaned with oxygen plasma). This ensured uniform coating and well dispersed fluorescent spot density for imaging in an area of 31.2 µm X 31.2 µm (a representative ROI in **figure S2A**). The resulting coated cover-slips were imaged in TIRF (movies were obtained till complete bleaching of the fluorescent spots). Individual RAβ oligomers bleached in a stepwise fashion. A few representative traces are shown in **figure S2B**.

The movies were analysed using Fiji (Image-J-win64, freely available image analysis program) for detecting the oligomers manually. The movies were thresholded and spots with intensity less than that were neglected. For a single spot, the diameter was restricted to 3×3 pixels (pixel size, 156 nm. To determine the stoichiometry, each spot was selected using the selection tool in Fiji and an intensity vs frames (i.e., time) plot was generated. The number of steps in which the fluorescence bleached in each spot was manually counted. This number gave the oligomer stoichiometry.

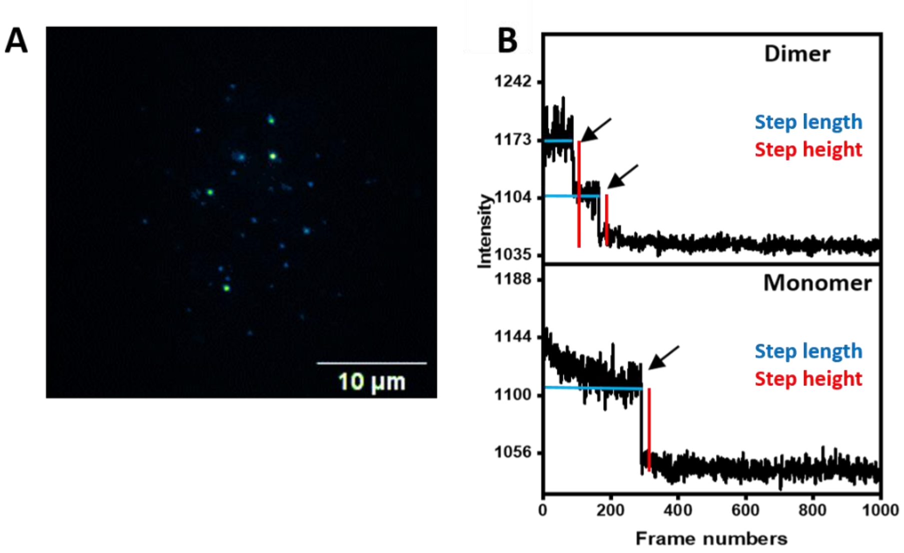

**Figure S2: Step-wise bleaching trajectories of RAβ oligomers. (A) A typical field of view of fluorescently labelled oligomers on a TIRF field of view. (B) Representative bleaching trajectories of a monomer and dimer. The step length and step heights are marked.**

For automated step detection of the oligomers a step-detection algorithm called pixel-classifier was developed in Napari (GitHub link provided below). In the software, once training of intensities (the fluorescent spots) and the background is done, it generates mask for each fluorescent spot and tracks it throughout the movie. Corresponding fitting of the intensity decay provides information on the step length, and step height from each fluorophore in an oligomer. Representative fitted traces is provided in figure S3.

Step detection was performed using a custom algorithm as described below:

1) Smoothening (Figure S5, B(red)):

   To remove spurious minor peaks, a 1D gaussian filter was applied on the particle intensity signal.

2) Normalisation (Figure S5, C(red)):

   The data was normalized and the first derivative was taken that converts the signal at the steps into peaks.

3) Peak detection (Figure S5, C):

   After normalising the signal, a threshold was applied to filter out smaller, spurious steps, resulting in the final positions of the peaks identifying the positions of the peaks.

4) Step information (Figure S5, D):

   Using the positions, we determine the length of the steps.

   Overlaying this information on the actual signal we determine the height of the steps.

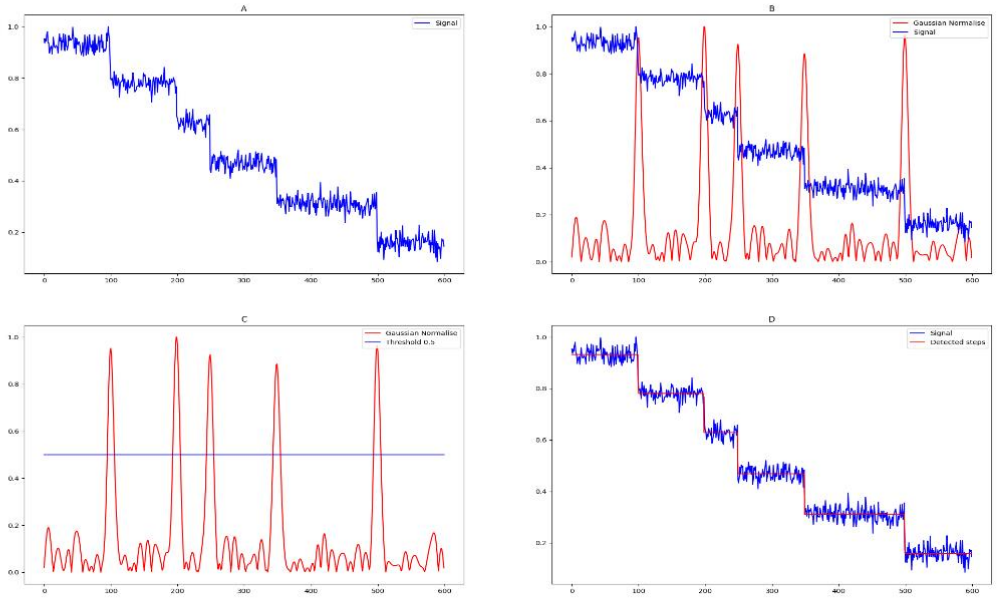

**Figure S3: A) Example stepwise intensity signal. B) First derivative signal overlaid on the actual signal where the sharp peaks coincides with step event in the original Signal. C) Threshold line to indicate the separation of the peaks. D) Step information overlayed on the original single.**

The GitHub link for the step detection algorithm is https://github.com/zeroth/step_detection.

**SI 4: Preparation of the supported lipid bilayer**

For measuring membrane affinity, membrane stoichiometry and QSLIP of RAβ oligomers, we prepared PPC 1:1:1 bilayer (POPC: POPG: Cholesterol in the molar ratio 1:1:1). The lipids 1-palmitoyl-2-oleoyl-sn-glycero-3-phosphocholine (16:0–18:1, PC) (POPC), 1-palmitoyl-2-oleoyl-sn-glycero-3-[phospho-rac-(1-glycerol)] (16:0–18:1 PG) (POPG) and cholesterol were purchased from Avanti Polar Lipids (Alabaster, AL) and from Sigma-Aldrich (St Louis, MO) respectively. SLBs were prepared using vesicle fusion method. Lipids were taken in appropriate amount and dissolved in chloroform to prepare a lipid film. The film was dried overnight in vacuum chamber. Lipid vesicle solution was prepared from the film using sonication. On a pre-cleaned coverslip chamber (prepared using cut PCR tubes and coverslip as the base), vesicle fusion was assisted using 10 mM $Ca^{2+}$. The entire system containing lipid vesicle solution, and calcium was left undisturbed in a water bath at 50°C. The prepared bilayer was incubated with 0.6 nM of RAβ oligomers for 30 minutes, washed with buffer, and finally imaged in the TIRF for membrane attachment.

**SI 5: Step lengths and QSLIP values of all the RAβ variants**

All the step length values (average and SEM) is listed in **table 1**. The values are in frame numbers (90 milliseconds per frame)

**Table 1:**

| | | Control | C-16 doxyl | C-5 doxyl | Tryptophan |
|---|---|---|---|---|---|
| **$RA\beta_{42}$** | **Monomer** | 75.65 ± 13.21 | 243.09 ± 65.26 | 86.64 ± 15.55 | 144.34 ± 19.89 |
| | **Dimer** | 50.82 ± 13.58 | 116.05 ± 25.09 | 63.40 ± 11.69 | 67.47 ± 13.71 |
| | | 154.23 ± 25.66 | 239.85 ± 25.86 | 148.33 ± 27.16 | 193.32 ± 25.79 |
| | **Trimer** | 24.11 ± 2.78 | 59.81 ± 6.20 | 54.91 ± 16.13 | 46.44 ± 8.08 |
| | | 56.81 ± 6.20 | 129.25 ± 11.74 | 104.69 ± 22.60 | 113.50 ± 16.17 |

| | | | | | |
|---|---|---|---|---|---|
| | | 128.29 ± 11.96 | 277.46 ± 32.40 | 186.08 ± 32.24 | 256.74 ± 31.50 |
| | **Tetramer** | 33.10 ± 4.22 | 44.54 ± 7.21 | 59.20 ± 28.45 | 46.57 ± 3.59 |
| | | 67.0 ± 12.04 | 94.72 ± 13.23 | 131.80 ± 49.23 | 107.57 ± 20.7 |
| | | 137.30 ± 18.83 | 194.27 ± 35.11 | 177.60 ± 63.27 | 181.21 ± 28.54 |
| | | 322.80 ± 52.97 | 347.63 ± 59.89 | 181.21 ± 28.54 | 277.78 ± 40.43 |
| **RAβ$_{40}$** | **Monomer** | 86.79 ± 12.64 | 101.64 ± 10.93 | 93.12 ± 18.80 | 73.39 ± 8.65 |
| | **Dimer** | 59.34 ± 9.89 | 50.0 ± 5.24 | 62.91 ± 14.19 | 60.59 ± 7.31 |
| | | 141.82 ± 16.65 | 125.35 ± 18.18 | 187.16 ± 42.68 | 156.83 ± 25.06 |
| | **Trimer** | 30.40 ± 3.10 | 30.81 ± 4.95 | 36.08 ± 7.23 | 32.61 ± 5.16 |
| | | 70.50 ± 7.70 | 72.67 ± 9.15 | 86.43 ± 18.27 | 80.61 ± 9.61 |
| | | 122.89 ± 12.09 | 147.81 ± 17.58 | 248.69 ± 77.80 | 164.16 ± 23.75 |
| | **Tetramer** | 14.67 ± 2.72 | 51.80 ± 10.70 | 35.0 ± 5.88 | 32.70 ± 4.20 |
| | | 32.67 ± 3.26 | 113.35 ± 31.50 | 79.60 ± 11.92 | 59.0 ± 7.51 |
| | | 57.50 ± 5.33 | 176.0 ± 38.96 | 122.75 ± 16.13 | 134.60 ± 39.7 |
| | | 215.5 ± 123.6 | 272.07 ± 30.17 | 199.0 ± 19.87 | 251.9 ± 98.7 |
| **RAβ$_{Cha}$** | **Monomer** | 89.40 ± 14.16 | 112.95 ± 27.09 | 104.9 ± 16.82 | 160.70 ± 29.07 |
| | **Dimer** | 70.53 ± 9.40 | 85.97 ± 22.92 | 53.56 ± 6.03 | 49.22 ±5.78 |
| | | 173.5 ± 24.67 | 206.08 ± 13.63 | 137.78 ± 10.98 | 125.92 ± 10.78 |
| | **Trimer** | 48.84 ± 6.64 | 51.14 ± 13.63 | 40.09 ± 4.30 | 59.09 ±10.52 |
| | | 105.53 ± 13.59 | 112.38 ± 21.65 | 90.20 ± 6.70 | 147.86 ± 24.47 |
| | | 193.69 ± 20.33 | 216.5 ± 27.90 | 202.60 ± 17.86 | 264.63 ± 37.32 |
| | **Tetramer** | 52.36 ± 10.85 | 39.77 ± 6.09 | 31.34 ± 2.52 | 35.42 ± 9.32 |
| | | 105.0 ± 23.22 | 87.77 ± 12.50 | 77.90 ± 12.78 | 84.71 ± 22.42 |
| | | 160.36 ± 32.39 | 171.67 ± 20.0 | 153.16 ± 19.63 | 175.42 ± 41.35 |
| | | 242.45 ± 39.54 | 273.33 ± 32.38 | 271.50 ± 34.48 | 255.0 ± 45.14 |

QSLIP values obtained from different oligomers of the RAβ isoforms are tabulated in table 2.

**Table 2:**

| | | C-16 doxyl | C-5 doxyl | Tryptophan |
|---|---|---|---|---|
| **RAβ$_{42}$** | **Monomer** | 3.21 ± 0.62 | 1.14 ± 0.40 | 1.73 ± 0.49 |
| | **Dimer** | 2.28 ± 0.31 | 1.24 ± 0.56 | 1.40 ± 0.58 |
| | | 1.55 ± 0.43 | 0.96 ± 0.34 | 1.38 ± 0.40 |
| | **Trimer** | 2.48 ± 0.60 | 2.27 ± 0.93 | 1.56 ± 0.52 |
| | | 2.28 ± 0.45 | 1.84 ± 0.59 | 1.74 ± 0.41 |
| | | 2.16 ± 0.45 | 1.45 ± 0.38 | 1.92 ± 0.39 |
| | **Tetramer** | 1.34 ± 0.38 | 1.78 ± 1.08 | 1.41 ± 0.28 |
| | | 1.41 ± 0.45 | 1.96 ± 1.08 | 1.61 ± 0.60 |
| | | 1.41 ± 0.43 | 1.29 ± 0.63 | 1.32 ± 0.39 |
| | | 0.86 ± 0.26 | 0.87 ± 0.43 | 0.86 ± 0.27 |
| **RAβ$_{40}$** | **Monomer** | 1.17 ± 0.29 | 1.07 ± 0.37 | 0.84 ± 0.22 |
| | **Dimer** | 0.84 ± 0.28 | 1.06 ± 0.42 | 1.02 ± 0.29 |
| | | 0.88 ± 0.23 | 1.32 ± 0.45 | 1.11 ± 0.31 |
| | **Trimer** | 1.01 ± 0.26 | 1.18 ± 0.35 | 1.07 ± 0.28 |
| | | 1.03 ± 0.24 | 1.23 ± 0.39 | 1.14 ± 0.26 |
| | | 1.20 ± 0.26 | 2.02 ± 0.83 | 1.34 ± 0.32 |
| | **Tetramer** | 3.53 ± 0.38 | 2.38 ± 0.84 | 2.22 ± 0.70 |
| | | 3.46 ± 0.31 | 2.44 ± 0.60 | 1.81 ± 0.41 |
| | | 3.06 ± 0.69 | 2.13 ± 0.47 | 2.34 ± 0.91 |
| | | 1.26 ± 0.28 | 0.92 ± 0.19 | 1.16 ± 0.26 |
| **RAβ$_{Cha}$** | **Monomer** | 1.26 ± 0.50 | 1.17 ± 0.37 | 1.80 ± 0.60 |
| | **Dimer** | 1.22 ± 0.48 | 0.76 ± 0.18 | 0.69 ± 0.17 |
| | | 1.19 ± 0.25 | 0.79 ± 0.18 | 0.72 ± 0.16 |
| | **Trimer** | 1.04 ± 0.42 | 0.82 ± 0.20 | 1.21 ± 0.37 |
| | | 1.06 ± 0.34 | 0.85 ± 0.17 | 1.40 ± 0.41 |
| | | 1.12 ± 0.26 | 1.04 ± 0.20 | 1.36 ± 0.33 |

| | | | | |
|---|---|---|---|---|
| | **Tetramer** | 0.76 ± 0.27 | 0.60 ± 0.17 | 0.67 ± 0.32 |
| | | 0.84 ± 0.30 | 0.74 ± 0.29 | 0.80 ± 0.39 |
| | | 1.07 ± 0.34 | 0.96 ± 0.32 | 1.09 ± 0.47 |
| | | 1.13 ± 0.32 | 1.12 ± 0.32 | 1.05 ± 0.35 |

**SI 6: Dose dependent increase in step length of RAβ42 oligomers**

The high QSLIP values of RAβ42 oligomers (monomer, dimer, and trimer) with C-16 doxyl suggested a very strong localization of its N-terminus in that region. However, except tetramer, none of the oligomers of RAβ40 showed localization near to the C-16 position. Thus, we wanted to check whether this difference is true. We made different preparations of PPC111 bilayer with two more concentrations, 1 mo l%, and 3 mol % of C-16 doxyl PC. With two such additional bilayers, we performed a dose dependence of step length for RAβ40 and RAβ42 oligomers. For comparison of the step lengths, we got statistically viable distribution till trimers. The results showed a dose dependent increase in the step length of RAβ42 oligomers, however RAβ40 oligomers showed no change in its step lengths (**figure S4**). This suggested that the high QSLIP value obtained from RAβ42 oligomers for C-16 doxyl indeed was because of a strong localization of its N-terminus near the carbon 16 position of the stearyl chain of the lipid, that is, deeper inside the bilayer.

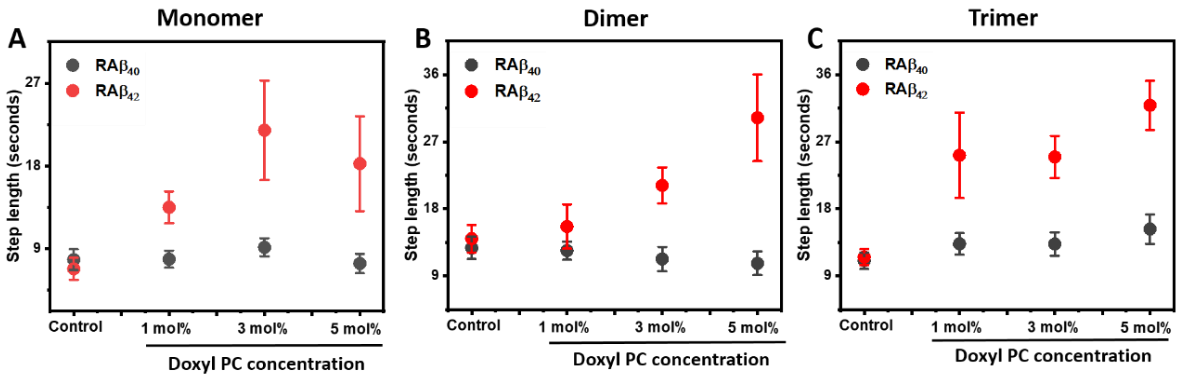

**Figure S4: Dose dependent increase in the step length values of RAβ42 oligomers as a function of the C-16 doxyl. This shows a N-terminus localization of the RAβ42 oligomers (till tetramer) near C-16 doxyl. However, the RAβ40 oligomers do not show any dose dependent increase in step length with C-16 doxyl. This suggests that till trimer, the N-terminus of RAβ40 do not penetrate the lipid bilayer as RAβ42 does.**

Thus, from the dose dependence measurements, we concluded that the N-terminus of Aβ42 oligomers, till trimer, could penetrate deep down the membrane (till C-16 doxyl). However, the oligomers of Aβ40 (except tetramer) were not able to penetrate deeper inside the bilayer (to the extent of RAβ42 oligomers). Thus membrane penetration could only be observed for both toxic RAβ40 and RAβ42.
